# Regulation of canonical Wnt signalling by the ciliopathy protein MKS1 and the E2 ubiquitin-conjugating enzyme UBE2E1

**DOI:** 10.1101/2020.01.08.897959

**Authors:** Katarzyna Szymanska, Karsten Boldt, Clare V. Logan, Matthew Adams, Philip A. Robinson, Marius Ueffing, Elton Zeqiraj, Gabrielle Wheway, Colin A. Johnson

## Abstract

A functional primary cilium is essential for normal and regulated signalling. Primary ciliary defects cause a group of developmental conditions known as ciliopathies, but the precise mechanisms of signal regulation by the cilium remain unclear. Previous studies have implicated the ubiquitin proteasome system (UPS) in regulation of Wnt signalling at the ciliary basal body. Here, we provide mechanistic insight into ciliary ubiquitin processing in cells and for a ciliopathy mouse model lacking the ciliary protein Mks1. *In vivo* loss of Mks1 sensitizes cells to proteasomal disruption, leading to abnormal accumulation of ubiquitinated proteins. To substantiate a direct link between MKS1 and the UPS, we identified UBE2E1, an E2 ubiquitin-conjugating enzyme that polyubiquitinates β-catenin, and RNF34, an E3 ligase, as novel interactants of MKS1. UBE2E1 and MKS1 colocalized, particularly during conditions of ciliary resorption, and loss of UBE2E1 recapitulates the ciliary and Wnt signalling phenotypes observed during loss of MKS1. Levels of UBE2E1 and MKS1 are co-dependent and UBE2E1 mediates both regulatory and degradative ubiquitination of MKS1. Furthermore, we demonstrate that processing of phosphorylated β-catenin occurs at the ciliary base through the functional interaction between UBE2E1 and MKS1. These observations suggest that correct β-catenin levels are tightly regulated at the primary cilium by a ciliary-specific E2 (UBE2E1) and a regulatory substrate-adaptor (MKS1), confirming the fundamental role of UPS defects in the molecular pathogenesis of ciliopathies.

## Introduction

Primary cilia are microtubule-based organelles that sense and transduce extracellular signals on many mammalian cells. The cilium has essential roles throughout development during mechanosensation ^1,2^, in transduction of multiple signalling pathways ^3-5^ and in the establishment of left-right asymmetry ^6^. Primary cilia have a complex ultrastructure with compartmentalization of molecular components that together form functional modules. Mutations in proteins that are structural or functional components of the primary cilium cause a group of human inherited developmental conditions known as ciliopathies ^7^. Examples of ciliopathies include Meckel-Gruber syndrome (MKS) and Joubert syndrome (JBTS). Many proteins that are mutated in ciliopathies, including the MKS1 protein ^8,9^, localize to the transition zone (TZ), a compartment of the proximal region of the cilium. Mutations in the *MKS1* gene cause about 15% of MKS, a lethal neurodevelopmental condition that is the most severe ciliopathy ^10^.

The MKS1 protein contains a B9/C2 domain with homologies to the C2 (calcium/lipid-binding) domain of the synaptotagmin-like and phospholipase families ^11^. MKS1 interacts with TMEM67, the transmembrane receptor encoded by the *TMEM67* gene ^12^, and two other B9/C2-domain containing proteins, B9D1 and B9D2 ^13^. B9D1, B9D2 and MKS1, are predicted to bind lipids in the ciliary membrane, and all three have been shown to localise at the ciliary TZ ^14^ forming components of a functional module (known as the “MKS-JBTS module”). This module contains other transmembrane proteins (TMEMs), namely the Tectonic proteins (TCTN1-3), TMEM17, TMEM67, TMEM231 and TMEM237, as well as other C2-domain proteins (jouberin, RPGRIP1L and CC2D2A) ^15-17^. TZ proteins are thought to form a diffusion barrier at the base of the cilium that restricts entrance and exit of both membrane and soluble proteins ^18^. The compartmentalization of the cilium is essential for the regulated translocation of signalling intermediates, most notably during Sonic hedgehog (Shh) signalling ^19^, and mutations of TZ components invariably cause Shh signalling defects during development ^20^. For example, mouse embryos from the *Mks1*^*Krc*^ knock-out mutant line have severe Shh signalling and left-right patterning defects during early embryonic development ^20^. Previously, we have described the *Mks1*^*tm1a(EUCOMM)Wtsi*^ knock-out mouse line, for which mutant embryos have de-regulated, increased canonical Wnt/β-catenin signalling and increased proliferation defects in the cerebellar vermis and kidney ^21^.

Other studies have shown that the ciliary apparatus restricts the activity of canonical Wnt/β-catenin signalling ^4,22,23^, although the mechanistic detail by which signal transduction is regulated remains unclear. One regulatory pathway involves the ciliary TZ protein jouberin (also known as AHI1), which shuttles β-catenin between the cytosol and nucleus in order to regulate Wnt signalling ^24^. However, ubiquitin-dependent proteasomal degradation by the ubiquitin-proteasome system (UPS) is the best-characterized mechanism for regulating canonical Wnt signalling ^25^. In the absence of a Wnt signal, cytoplasmic β-catenin is phosphorylated in a complex of proteins (referred to as the destruction complex) that include axin, adenomatous polyposis coli (APC), and glycogen synthase kinase 3 (GSK-3)^26-28^. Subsequent ubiquitination of β-catenin leads to its degradation by the proteasome, meaning that in the absence of Wnt signalling the steady state levels of cytoplasmic β-catenin are low. Part of this regulation appears to be mediated by a functional association of the ciliary apparatus with the UPS ^29^, and UPS components have been shown to interact with ciliopathy proteins (e.g. USP9X and lebercilin) ^30^. RPGRIP1L (a ciliary TZ protein mutated in a range of ciliopathies including MKS and JBTS) has been reported to interact with the proteasome proteins, PSMD3 and PSMD5 ^31^. Furthermore, discrete localization of ubiquitin has been observed at the ciliary base suggesting that UPS processing can be constrained and regulated by the cilium ^31^. However, the mechanistic basis to substantiate the association between the UPS and ciliary apparatus remains unclear and, in particular, it is unknown if the pathomechanism of Wnt signalling defects in ciliopathies depends on defective regulation of β-catenin localization and processing by ciliary proteins.

Here, we describe the interaction and functional association of MKS1 with ciliary UPS components, specifically the E2 ubiquitin-conjugating enzyme UBE2E1 (also known as UbcH6) and the E3 ubiquitin ligating enzyme RNF34. In addition to ciliogenesis defects, loss of MKS1 causes deregulation of both proteasome activity and canonical Wnt/β-catenin signalling. These cellular phenotypes are also observed after loss of UBE2E1. MKS1 and UBE2E1 colocalize during conditions of cilia resorption, and levels of MKS1 and UBE2E1 are co-dependent. We show that in the absence of MKS1, levels of ubiquitinated proteins, including β-catenin, are increased. Furthermore, polyubiquitination of MKS1 is dependent on both UBE2E1 and RNF34, and lysine (Lys)63-linked polyubiquitination of MKS1 is dependent on UBE2E1. This suggests that regulation of intracellular signalling, specifically canonical Wnt/β-catenin signalling, can be regulated and constrained at the primary cilium by a ciliary-specific E2 and MKS1, a substrate-adaptor.

## Results

### Mks1 mutation causes deregulation of proteasome activity

Loss of ciliary basal body proteins perturbs both UPS function and Wnt signalling ^29^, and we have previously reported de-regulated increases of canonical Wnt signalling in *Mks1*^*-/-*^ mutant mice ^21^. To investigate the mechanistic basis for regulation of canonical Wnt/β-catenin signalling and possible UPS processing of β-catenin by a ciliary protein, we first characterized these processes in cells and tissues lacking functional MKS1. We derived immortalized dermal fibroblasts from a human MKS patient, carrying compound heterozygous *MKS1* mutations [c.472C>T]+[IVS15-7_35del29] causing the predicted nonsense and splice-site null mutations [p.R158*]+[p.P470*fs*\*562] ^10^ (**Supplementary Figure 1a**) leading to loss of MKS1 protein (**Supplementary figure 1b-c**). *MKS1*-mutated fibroblasts had decreased cilia incidence and length (**Supplementary figure 1d**), and de-regulated canonical Wnt/β-catenin signalling (**Figure 1a**). *MKS1*-mutated fibroblasts had moderately increased levels of total β-catenin and the Wnt downstream target cyclin D1 (**Figure 1a**). SUPER-TOPFlash reporter assays confirmed that increased levels of β-catenin in *MKS1*-mutated fibroblasts caused de-regulated increases in canonical Wnt signalling in response to Wnt3a (a canonical Wnt ligand; **Figure 1b**). Treatment with the non-specific proteasome inhibitor MG-132 also increased levels of phosphorylated β-catenin (**Figure 1a**). Since β-catenin is phosphorylated to mark it for processing by the 26S proteasome, we also tested if proteasome enzymatic activity was affected in *MKS1*-mutated fibroblasts. We observed increased proteasome activity, which was inhibited by treatment with lactacystin that targets the 20S catalytic core of the proteasome, as well as moderate increased levels of the proteasome subunit α7 (**Figure 1c**).

**Figure 1:**
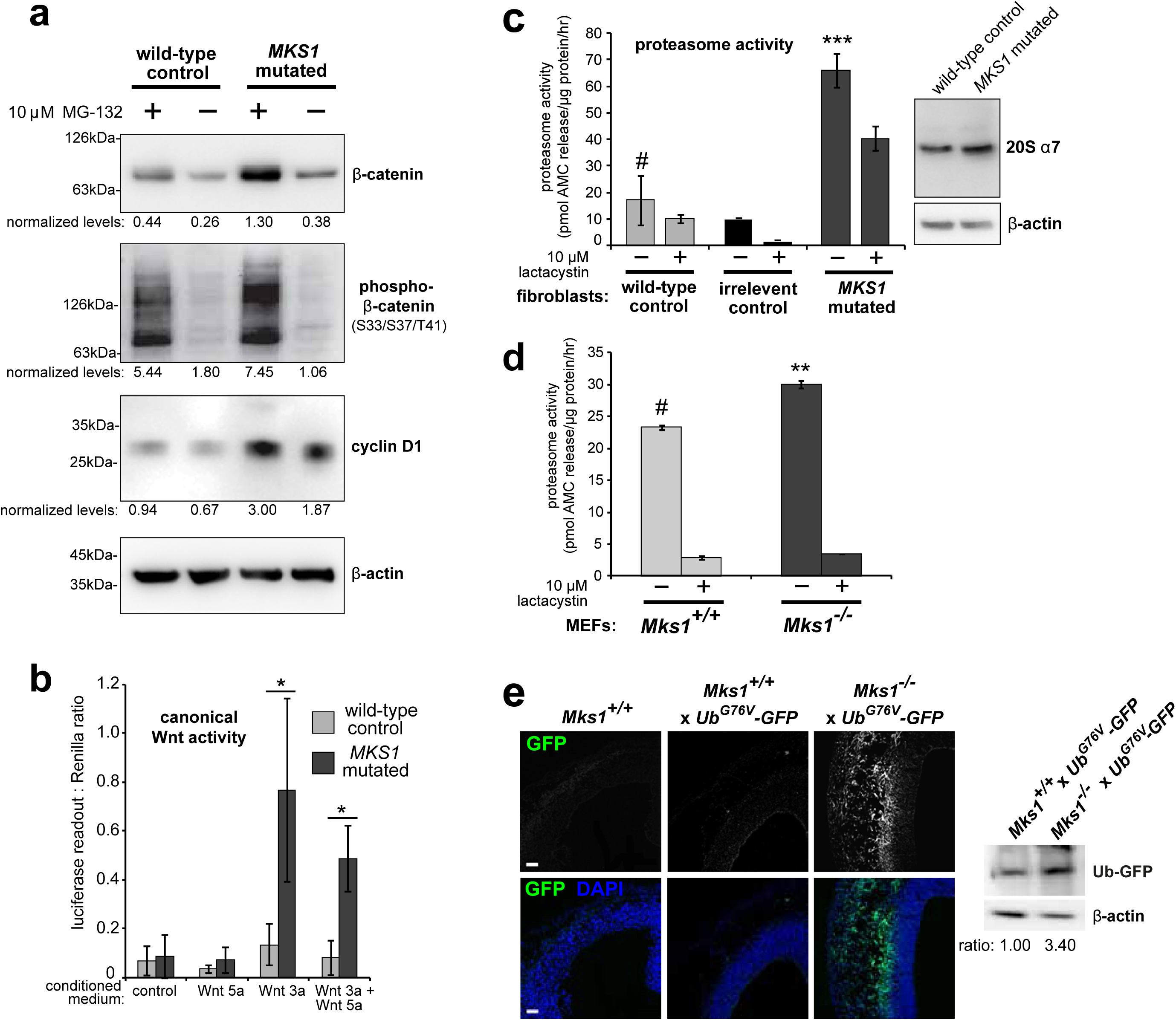
Deregulation of canonical Wnt signalling and proteasome activity following loss or mutation of MKS1. **(a)** Immunoblots for total soluble β-catenin, phospho-β-catenin, cyclin D1 and β-actin (loading control) in either wild-type normal or *MKS1*-mutated immortalized human fibroblasts from an MKS patient (MKS-562) following treatment with MG-132 proteasome inhibitor (+) or vehicle control (-). **(b)** SUPER-TOPFlash assays of canonical Wnt signalling activity in human *MKS1*-mutated fibroblasts compared to wild-type control fibroblasts following treatment with control conditioned medium, Wnt5a, Wnt3a, or a mixture of Wnt3a and Wnt5a media, as indicated. Statistical significance of pairwise comparisons is shown (* indicates *p* < 0.05, paired two-tailed Student t-test). Error bars indicate s.e.m. with results shown for four independent biological replicates. **(c)** Proteasome activity assays for wild-type or *MKS1*-mutated human fibroblasts from patient MKS-562 or an irrelevant control (*ASPM-*mutant fibroblasts), following treatment with c-lactacystin-β-lactone (+) or vehicle control (-). Statistical significance of pairwise comparison as for (b); *** indicates *p* < 0.001 for three independent biological replicates. Immunoblots show levels of the 20S proteasome α7 subunit compared to β-actin loading control. **(d)** Protease activity assays of crude proteasome preparations from *Mks1*^*+/+*^ or *Mks1*^*-/-*^ mouse embryonic fibroblasts (MEFs), expressed as pmol AMC released per μg proteasome per hr. Treatment with lactacystin is the assay control. Statistical analysis as for (b); ** indicates *p* < 0.01 for three independent biological replicates. **(e)** Proteasome defects and accumulation of GFP-tagged ubiquitin (GFP; green) in *Mks1*^*-/-*^ x *Ub*^*G76V*^*-GFP* E12.5 embryonic cerebral neocortex. Immunoblot for GFP in *Mks1*^*-/-*^ x *Ub*^*G76V*^*-GFP* and wild-type littermate E12.5 embryo protein lysates, with immunoblotting for β-actin as a loading control, showing accumulation of GFP-tagged ubiquitin (Ub-GFP) in *Mks1*^*-/-*^.

To substantiate an *in vivo* association between de-regulated canonical Wnt signalling and proteasome activity in the ciliopathy disease state, we crossed the *Mks1*^*tm1a(EUCOMM)Wtsi*^ knock-out mouse line ^21^ with the *Ub*^*G76V*^*-GFP* transgenic reporter line. *Ub*^*G76V*^*-GFP* constitutively degrades transgenic ubiquitinated-GFP, leading to an absence of GFP signal if proteasome processing is unimpaired ^32^. Confirming our observations with human *MKS1*-mutated fibroblasts, *Mks1*^*-/-*^ x *Ub*^*G76V*^*-GFP* mouse embryonic fibroblasts (MEFs) also had de-regulated proteasome enzymatic activity (**Figure 1d**) compared to *Mks1*^*+/+*^ x *Ub*^*G76V*^*-GFP* wild-type littermate MEFs. Furthermore, *Mks1*^*-/-*^ x *Ub*^*G76V*^*-GFP* mutant embryos at embryonic day E12.5 had increased levels of GFP, detected by both epifluorescence confocal microscopy and western blotting, in the neocortex (**Figure 1e**) and other tissues (**Supplementary Figure 2a-b**) compared to wild-type littermate controls. Autofluorescence was virtually undetectable in these tissues (data not shown). The accumulation of GFP in mutant embryonic tissues indicated that constitutively expressed GFP-tagged ubiquitin was not properly degraded, suggesting that these tissues harboured proteasomal defects. These defects accompanied an absence of ciliogenesis and increased levels of active β-catenin in the neuroepithelium of *Mks1*^*-/-*^ x *Ub*^*G76V*^*-GFP* mutant ventricular zone (**Supplementary Figure 2b**).

### MKS1 interacts with the E2 ubiquitin-conjugation enzyme UBE2E1, with colocalization during cilia resorption

To understand why mutation or loss of MKS1 causes de-regulated increases of both proteasome activity and canonical Wnt/β-catenin signalling, we sought to identify MKS1-interacting proteins. We performed a yeast two-hybrid screen using amino acids 144-470 of MKS1 that contain the B9/C2 domain as bait (**Figure 2a**) and identified the E2 ubiquitin conjugation enzyme UBE2E1 (also known as UbcH6) ^33^ as an interactant of MKS1 (**Figure 2b**). We confirmed this interaction by a “one-to-one” yeast two-hybrid assay (**Figure 2c**). Additionally, we identified the E3 ubiquitin ligase RNF34 and confirmed its interaction and colocalization with MKS1 (**Figure 2b, Supplementary Figure 3a-b**). In support of a possible role of MKS1 in regulating ubiquitinated signalling proteins, UBE2E1 has been described to function as an E2 with the E3 JADE-1 during the ubiquitination of β-catenin ^34^. We therefore further substantiated the interaction of UBE2E1 with MKS1. We purified GST-tagged UBE2E1 and confirmed the interaction between MKS1 and UBE2E1 by a GST pull-down assay (**Figure 2d**). The interaction between endogenous MKS1 and UBE2E1 was confirmed by co-immunoprecipitations (co-IPs) using anti-MKS1 (**Figure 2e**). This was further corroborated when an interaction between endogenous UBE2E1 and exogenously expressed cmyc-tagged MKS1 was detected by co-IP with an anti-UBE2E1 antibody (**Figure 2f**).

**Figure 2:**
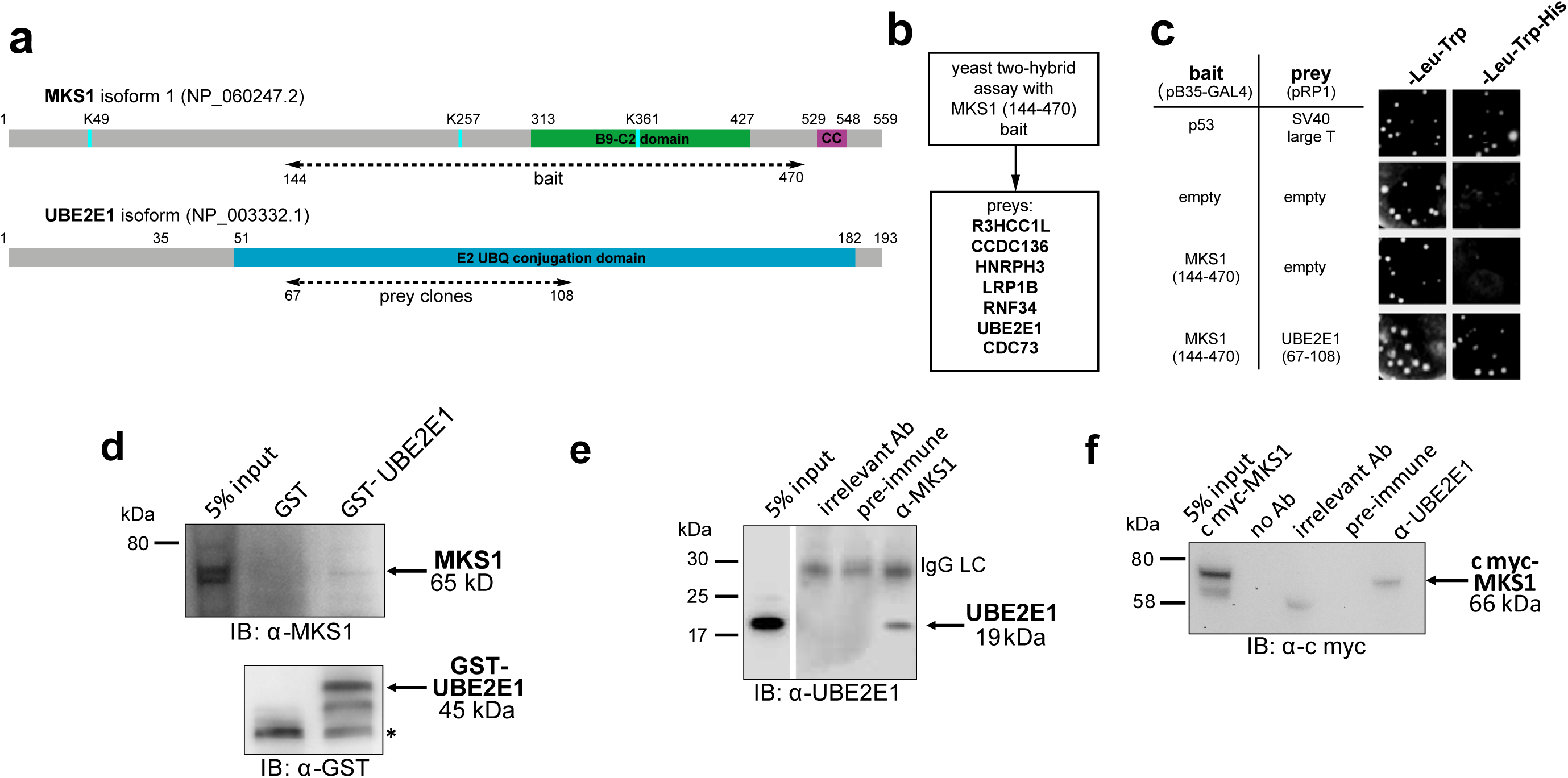
The E2 ubiquitin conjugation enzyme UBE2E1 interacts with MKS1. **(a)** Domain structure of MKS1 and UBE2E1 proteins for the indicated isoform showing the locations of the B9/C2 domain, putative ubiquitinated lysines in blue (predicted by UbPred), a predicted coiled-coil (CC) motif, and the E2 ubiquitin (UBQ) conjugation domain in UBE2E1. Numbering indicates the amino acid residue. Dashed lines indicate the region used as “bait” in MKS1 for the yeast two-hybrid assay and the “prey” clones in the UBE2E1 interactant. **(b)** List of preys identified in the MKS1 Y2H screen **(c)** Left panel: yeast “one-to-one” assays for the indicated bait, prey and control constructs. Right panel: only colonies for the positive control (p53+SV40 large T) and MKS1 bait+UBE2E1 prey grew on triple dropout (-Leu -Trp -His) medium. **(d)** GST pulldown of endogenous MKS1 by GST-UBE2E1 fusion protein but not GST. **(e)** Co-immunoprecipitation (co-IP) of endogenous UBE2E1 by rabbit polyclonal anti-MKS1, but not pre-immune serum or an irrelevant antibody (Ab; anti cmyc); IgG light chain (LC) is indicated. **(f)** Co-IP of exogenously expressed cmyc-MKS1 by anti-UBE2E1 but not pre-immune serum or an irrelevant antibody.

### The UBE2E1-MKS1 interaction is required for cilia resorption

Having confirmed UBE2E1-MKS1 interaction *in vitro* and in cells, we wondered if this was important for cilia function. UBE2E1 and MKS1 co-localized at the basal body in a subset of confluent, ciliated hTERT-immortalized retinal pigment epithelium RPE1 and ARPE19 cells during G_0_ of the cell cycle following serum starvation for 48hr ^35^ (**Figure 3a-d**). Serum starvation, followed by re-addition of serum for 3hr, caused rapid cilia resorption ^35^ with further significant colocalization of UBE2E1 and MKS1 at the basal body (**Figure 3b** and **3d**). This suggests that the interaction between MKS1 and UBE2E1 is particularly important during the process of cilia resorption. Furthermore, UBE2E1 co-localised with γ-tubulin at the basal body in *MKS1*-mutated fibroblasts, unlike in normal wild-type cells (data not shown). This suggests that in normal cell conditions MKS1 excludes UBE2E1 from basal body localisation.

**Figure 3:**
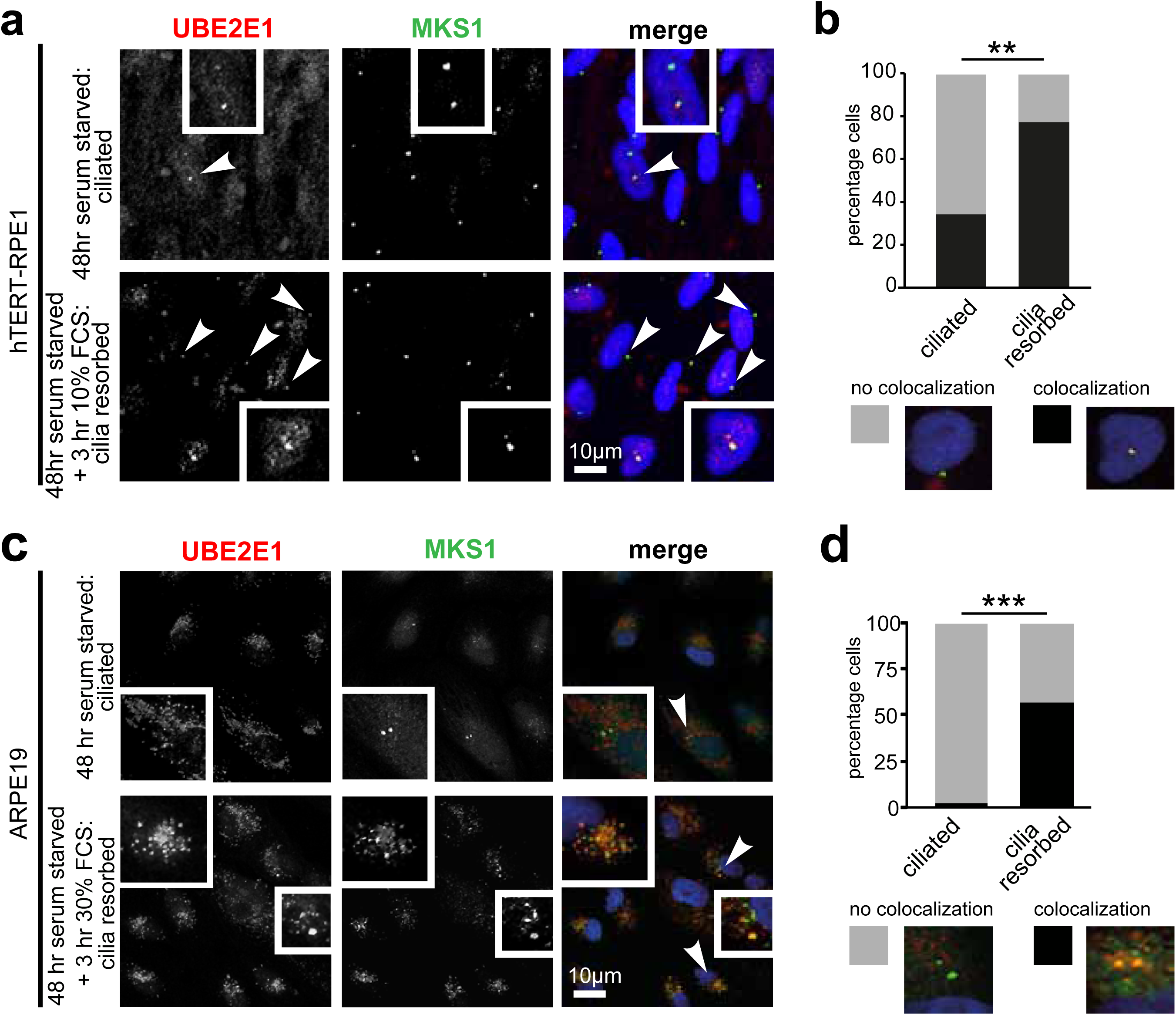
Co-localisation of MKS1 and UBE2E1 under conditions of ciliary resorbtion. **(a)** MKS1 (green) and UBE2E1 (red) partially colocalize at the basal body/centrosome in human wild-type hTERT-RPE1 cells, particularly when induced to resorb cilia by treatment with 10% FCS after 72 hr of serum starvation. White arrowheads indicate cells magnified in insets. Scale bar = 10 μm. **(b)** Bar graph indicates the percentage of cells in which MKS1 and UBE2E1 co-localize at the basal body (black), and the percentage without co-localization (grey) for three independent biological replicates, with examples shown of representative cells. **(c)** Figure details as for (a) showing partial co-localization of MKS1 and UBE2E1 in human ARPE19 cells. **(d)** Bar graph details as for (b). Data in (b) and (d) was analysed by two-way ANOVA followed by Tukey’s multiple comparison test (statistical significance of comparisons indicated by ** *p*<0.01, *** *p*<0.001).

### UBE2E1 mutation or loss causes ciliogenesis defects, and de-regulated increases in both proteasome activity and Wnt/β-catenin signalling

Since correct UPS function appears to be required for ciliogenesis ^31,40,41^, we next asked if loss or mutation of UBE2E1 had an effect on ciliogenesis. UBE2E1 is an enzyme that transfers ubiquitin to a substrate, with or without the presence of an E3, in a reaction that is dependent on an active enzymatic domain. To assess if enzymatic activity of UBE2E1 is necessary for correct ciliogenesis, we mutated the active site cysteine residue 131 to serine ^36^ to make a dominant negative (DN) catalytically-inactive form of UBE2E1. Over-expression of the Cys131Ser form of UBE2E1 caused significant loss and shortening of cilia in mouse inner medullary collecting duct (mIMCD3) cells (**Figure 4a-b**), suggesting that catalytically active UBE2E1 is required for normal ciliogenesis. Over-expression of wild-type (WT) UBE2E1 had a moderate dominant negative effect on cilia length only (**Figure 4a-b**).

**Figure 4:**
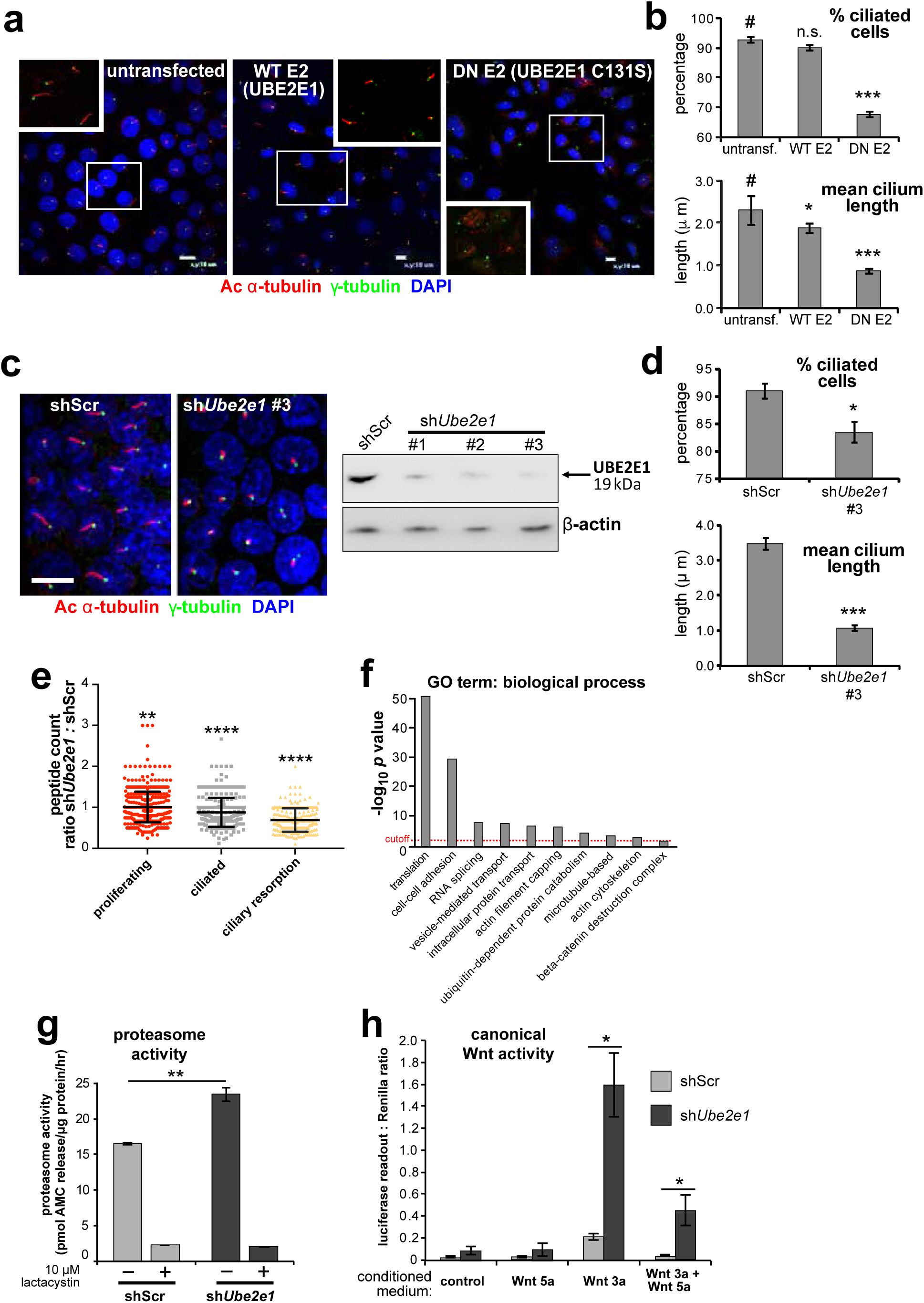
UBE2E1 is required for regulation of ciliogenesis, proteasome activity and canonical Wnt signalling. **(a)** Primary cilia in mIMCD3 cells following transfection with either wild-type (WT) UBE2E1 (E2) or dominant negative (DN) UBE2E1 carrying the active site mutation C131S, compared to mock-transfected negative control. Scale bars = 10μm. **(b)** For experiments shown in (a), statistical significance of pairwise comparisons with control (#) for three independent biological replicates are shown (n.s. not significant, * *p* < 0.05, ** *p*<0.01, *** *p* < 0.001; unpaired two-tailed Student t-test; error bars indicate s.e.m.). **(c)** shRNA-mediated knockdown of *Ube2e1* in stably-transfected mIMCD3 cell-line #3 causes decreased ciliary incidence and length. Scale bar = 10μm. Immunoblot shows loss of UBE2E1 protein expression compared to β-actin loading control following shRNA knockdown. **(d)** Bar graphs quantifying decreased ciliary incidence and length as for (b). **(e)** Scatter plot of relative differences in the proteins pulled-down by anti-MKS1 immunoprecipitations, under different conditions of ciliogenesis (proliferating cells, ciliated cells, cells undergoing ciliary resorption), expressed as the ratios of peptide counts for shScr: sh*Ube2e1* knockdowns. Statistical significance of pairwise comparisons for each set of ratios was calculated as for (b) (paired two-tailed Student t-tests). Error bars indicate s.d. Full data-sets are available in **Supplementary Dataset 1. (f)** Bar graph of -log_10_ *p* values for significantly enriched GO terms (biological processes) for proteins included in (e), with cut-off for *p*<0.05 indicated by the red dashed line. Enrichment for GO terms was analyzed by using DAVID (https://david.ncifcrf.gov/). **(g)** Protease activity assays of crude proteasome preparations from shScr and sh*Ube2e1* mIMCD3 knockdown cells, showing increased proteasomal activity in sh*Ube2e1* as assayed by pmol AMC released per μg proteasome per hour. Treatment with lactacystin is the assay control. Statistical significance of pairwise comparisons as for (b). **(h)** SUPER-TOPFlash assays of canonical Wnt signalling activity in sh*Ube2e1* cells compared to shScr following treatment with control conditioned medium, Wnt5a, Wnt3a, or a mixture of Wnt3a and Wnt5a media, as indicated. Statistical significance of pairwise comparisons of at least four independently replicated experiments as for (b).

To model the effect of UBE2E1 loss on ciliogenesis, we first used pooled and individual siRNA duplexes targeting *Ube2e1* in mIMCD3 cells. This affected ciliogenesis in mIMCD3 cells by reducing cilia incidence and length, but achieved only moderate knockdown of UBE2E1 protein levels (**Supplementary Figure 4a**). To ensure more robust, long-term knockdown of UBE2E1, we derived stably-transfected mIMCD3 cell-lines with three different *Ube2e1* shRNA constructs. Each *Ube2e1* shRNA construct reduced UBE2E1 protein levels (compared to cells expressing scrambled shRNA), and significantly reduced both numbers of ciliated cells and mean cilium length (**Figure 4c-d**). To understand the effect loss of Ube2e1 had on Mks1, we pulled down Mks1 at different ciliogenesis stages in control cells and cells with stable knockdown of *Ube2e1* followed by identification of interacting proteins by LC-MS/MS mass spectrometry analysis (**Supplementary Dataset 1**). We observed significant decreases in peptide counts for sh*Ube2e1* knockdown cells across different conditions of ciliogenesis (**Supplementary Dataset 1**; χ^2^ test *p*<0.05), particularly under conditions of ciliary resorption (**Figure 4e**). Analysis for enrichment of GO terms identified specific biological processes that included “cell-cell adhesion”, “ubiquitin-dependent protein catabolism”, “actin filament capping” and “beta-catenin destruction complex” (**Figure 4f**). Interestingly, the “actin filament capping” term includes known interactants of ciliopathy proteins such as filamin A (Flna) ^37^ (**Supplementary Figure 4b**).

Since UBE2E1 and MKS1 both interact and co-localize, we next determined if UBE2E1 loss reiterates the cellular phenotypes caused by MKS1 loss or mutation. Indeed, we observed increased proteasome enzymatic activity compared to scrambled shRNA (shScr) negative control cells (**Figure 4g**). Furthermore, in agreement with *MKS1*-mutated fibroblasts and *Mks1*^*-/-*^ MEFs, sh*Ube2e1* knock-down cells had de-regulated canonical Wnt/β-catenin signalling in response to Wnt3a (**Figure 4h**). These data highlight a possible important role of UBE2E1 in mediating correct protein ubiquitination, proteasome function and Wnt signalling in the context of cilia. The striking similarities in ciliary phenotypes suggest a close functional association between MKS1 and UBE2E1, and led us to hypothesize that they are placed in the same signalling pathway.

### Mutual inhibition of MKS1 and UBE2E1 protein levels

To further investigate the functional association between MKS1 and UBE2E1, we tested if de-regulated Wnt signalling could be rescued by over-expression experiments. In a control experiment, expression of cmyc-tagged MKS1 partially rescued normal canonical Wnt signalling responses to Wnt3a in *MKS1*-mutated fibroblasts (**Figure 5a**). However, expression of FLAG-tagged UBE2E1 led to almost complete rescue of normal Wnt signalling responses (**Figure 5a**). Conversely, expression of MKS1 in sh*Ube2e1* knock-down cells also rescued canonical Wnt signalling (**Figure 5b**), suggesting co-dependency between MKS1 and UBE2E1. We confirmed this following transient siRNA knockdown of MKS1 that caused significantly increased levels of UBE2E1 in cells (**Figure 5c**). In the reciprocal experiment, MKS1 protein levels were significantly increased in sh*Ube2e1* knock-down cells (**Figure 5c**). To further support co-dependency, we over-expressed both MKS1 and UBE2E1. At higher levels of UBE2E1, we observed a moderate decrease in MKS1 levels (**Figure 5d**). Conversely, expression of high levels of MKS1 caused a significant decrease in UBE2E1 protein levels (**Figure 5d**). These results show a striking co-dependency in protein levels between MKS1 and UBE2E1, suggesting inhibitory roles for each of these proteins on the protein level of the other.

**Figure 5:**
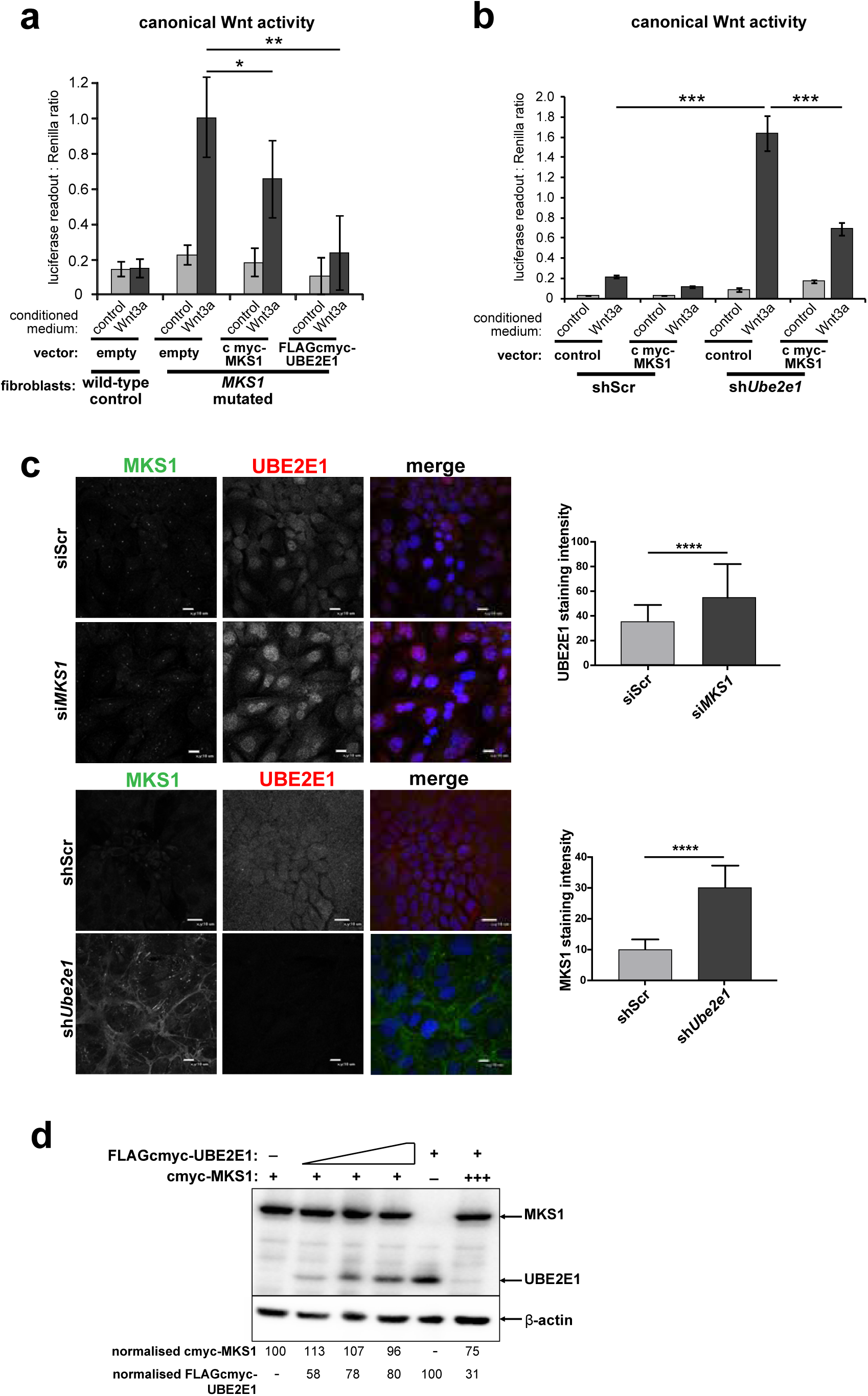
Co-dependant regulation of MKS1 and UBE2E1. **(a)** SUPER-TOPFlash assays in wild-type or *MKS1*-mutated fibroblasts, following transient co-transfection with either exogenous control, MKS1-cmyc or UBE2E1-FLAG-cmyc, and treatment with either Wnt3a or control conditioned medium. Statistical significance of the indicated pairwise comparisons with control for three independent biological replicates are shown (* *p* < 0.05, ** *p*<0.01, *** *p* < 0.001, **** *p* < 0.0001; unpaired two-tailed Student t-test; error bars indicate s.e.m.) **(b)** SUPER-TOPFlash assays in shScr and sh*Ube2e1* cell-lines, following transient co-transfection with either exogenous cmyc-MKS1 or empty plasmid construct (control) and treatment with either Wnt3a or control conditioned medium, as indicated. Statistical comparisons as for (a). **(c)** Top panel: increased per cell staining intensity for UBE2E1 following *MKS1* siRNA knockdown Bottom panel: increased per cell staining intensity for MKS1 in *Ube2e1* mIMCD3 knockdown cells Scale bars= 10μm. Bar graphs quantitate staining intensities for three independent biological replicates. Statistical significance of pairwise comparisons as for (a). **(d)** HEK293 cells were transiently transfected with control vector (-), constant (+) or high (+++) levels of cmyc-MKS1, and/or increasing levels of FLAG-cmyc-UBE2E1. Levels were normalized to β-actin loading control. MKS1 levels decreased with increasing levels of UBE2E1, whereas high levels of MKS1 caused loss of UBE2E1.

### MKS1 is polyubiquitinated and its polyubiquitination depends on UBE2E1

UBE2E1 is an E2 ubiquitin conjugating enzyme, and we next tested the obvious hypothesis that it participates in polyubiquitination and targeting MKS1 for degradation. We therefore investigated if MKS1 is indeed tagged with ubiquitin chains and if absence of UBE2E1 affects ubiquitination of MKS1. We determined MKS1 levels and its polyubiquitination status in different ciliogenesis conditions, namely: proliferating cells (grown in normal medium supplemented with serum); ciliated cells (quiescent cells grown in serum-deprived medium); and cells undergoing ciliary resorption (grown in serum-deprived medium, followed by serum re-addition for 3hr). sh*Ube2e1* knock-down cells consistently had significantly increased levels of MKS1 as well as polyubiquitinated MKS1 (**Figure 6a**; *p*<0.05 paired Student t-test between shScr and sh*Ube2e1*). The highest levels of MKS1 were observed in cells undergoing cilia resorption, when MKS1 and UBE2E1 co-localisation is the strongest. Furthermore, expression of exogenous UBE2E1 led to moderate decreases in MKS1 levels for both shScr and sh*Ube2e1* knock-down cells, suggesting that UBE2E1 inhibits both MKS1 levels and polyubiquitination of this protein.

**Figure 6:**
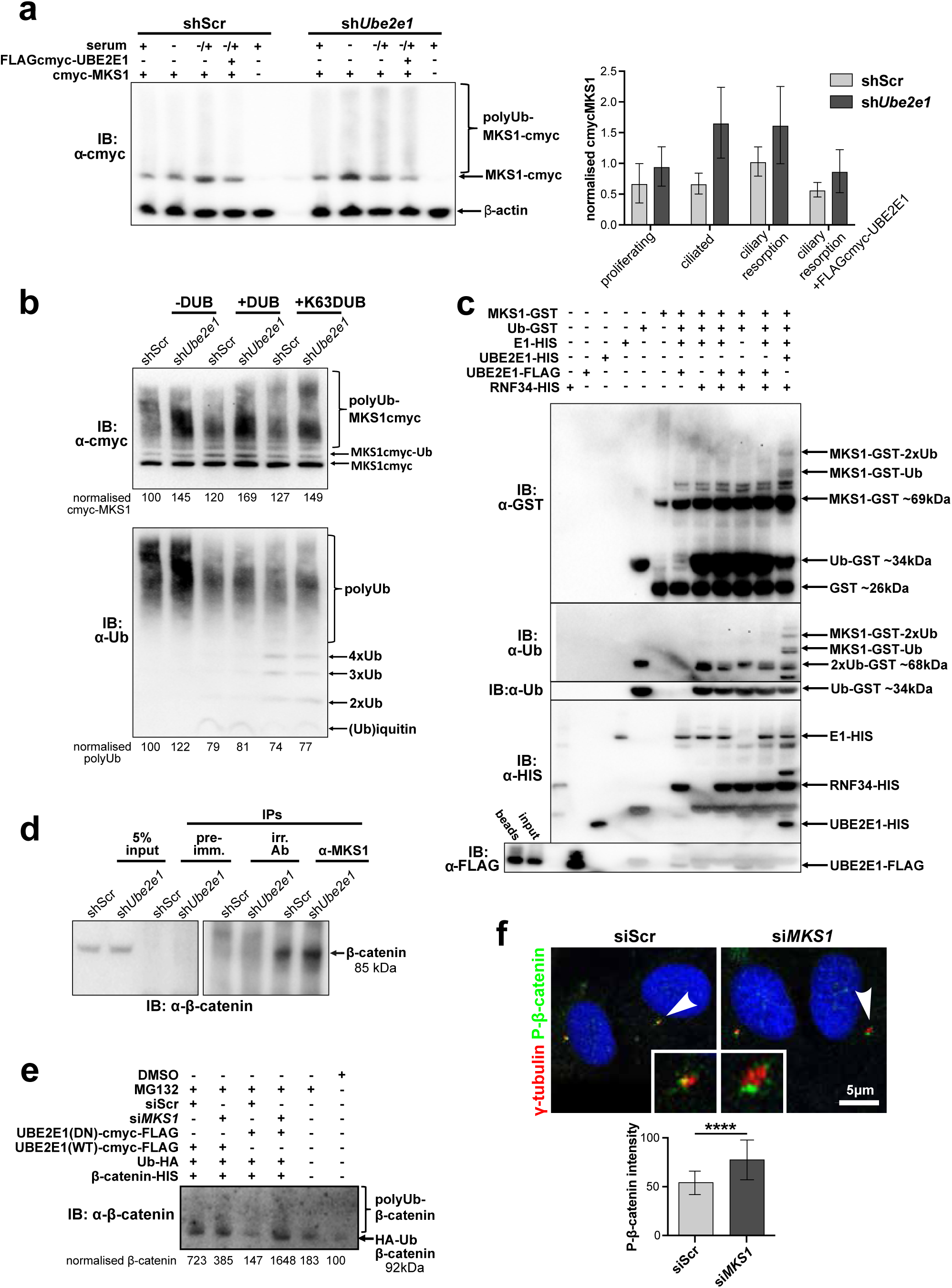
MKS1 is ubiquitinated and its ubiquitynation depends on UBE2E1. **(a)** shScr and sh*Ube2e1* mIMCD3 knockdown cells transiently transfected with cmyc-MKS1 and/or FLAG-cmyc-UBE2E1 under different conditions of ciliogenesis: proliferating cells grown in media containing serum (+); ciliated cells grown in the absence of serum (-); and cells undergoing ciliary resorption grown in the absence of serum followed by 2hr incubation in media with serum (-/+). Increased levels of cmyc-MKS1 and smears representing poly-ubiquitinated (polyUb) cmyc-MKS1 in sh*Ube2e1* cells are indicated. Addition of exogenous FLAG-cmyc-UBE2E1 partially rescued correct MKS1 levels and ubiquitination. Bar graph quantitates cmyc-MKS1 levels normalized to β-actin levels for three independent biological replicates. Data was analysed by two-way ANOVA followed by Tukey’s multiple comparison test (statistical significance of comparison between shScr and sh*Ube2e1* is *p*<0.05). **(b)** TUBE experiment confirming ubiquitination of cmyc-MKS1. Consistently increased levels of polyubiquitinated cmyc-MKS1 were observed in sh*Ube2e1* knockdown cells. Broad-range deubiquitinating enzymes (+DUB) and K63-specific (+K63DUB) deubiquitinating enzyme were used to assess the type of MKS1 ubiquitination. Normalised band intensities are shown below the blots. **(c)** *In vitro* ubiquitination assay for MKS1-GST fusion protein, for the indicated controls and reaction conditions, showing the addition of one (Ub) and two (2xUb) ubiquitins in the presence of both UBE2E1 and RNF34. **(d)** Immunoblot showing increased co-immunoprecipitation of β-catenin by anti-MKS1 in sh*Ube2e1* knockdown cells compared to shScr control cells. **(e)** TUBE pulldown followed by β-catenin immunoblotting, showing polyubiquitination of β-catenin was increased following *MKS1* knockdown and in the presence of the dominant negative (DN) catalytically-inactive form of UBE2E1 compared to wild-type (WT) UBE2E1. **(f)** Immunofluorescence staining of hTERT-RPE1 cells showing co-localization of phosphorylated (P)-β-catenin (green) with γ-tubulin (red) at the base of cilia (arrowheads). P-β-catenin staining intensity was measured using ImageJ, showing a significant increase following *MKS1* knockdown (paired two-tailed Student t-test, **** *p* < 0.0001 for three independent biological replicates; >40 cells quantified per replicate). Scale bar=5 μm.

To substantiate the central role of UBE2E1 in regulating MKS1 levels, we confirmed that sh*Ube2e1* cells had increased levels of polyubiquitinated cmyc-tagged MKS1 using TUBE (Tandem Ubiquitin Entity) assays. Total polyubiquitinated proteins from cell extracts were pulled-down using TUBEs bound to agarose beads, resolved by SDS-PAGE and analysed by western blotting using an anti-cmyc antibody. TUBE assays confirmed that MKS1 was polyubiquitinated (**Figure 6b**, upper panel). Treatment of pull-downs with either a pan-specific deubiquitinase (DUB) and a DUB specific for Lys63-linked polyubiquitination confirmed that MKS1 polyubiquitination occurred through both Lys63 and other ubiquitin lysine linkages such as Lys48 (**Figure 6b**, lower panel). Importantly, these results suggest that ubiquitination of MKS1 has dual functions in targeting this protein for degradation, as well as other regulatory functions through Lys63. Although UBE2E1 could be an E2 in an MKS1 degradation pathway, our data showed that loss of UBE2E1 caused an increase in the levels of polyubiquitinated MKS1 consistent with an inhibitory function for UBE2E1 in ubiquitinating MKS1. To test this alternative hypothesis, we therefore performed *in vitro* ubiquitination assays in which purified MKS1 was used as a substrate of the reaction, purified UBE2E1 was the E2 and RNF34 was a possible cognate E3. We observed mono- and di-ubiquitination of MKS1 by UBE2E1 in the presence of RNF34 (**Figure 6c**), confirming the role of UBE2E1 in MKS1 ubiquitination.

### MKS1 and UBE2E1 interact to regulate β-catenin ubiquitination

Monoubiquitination by UBE2E1 has been previously described ^38^ and UBE2E1 has also been shown to be an E2 ubiquitin-conjugating enzyme required for β-catenin polyubiquitination ^34^. These studies suggest that UBE2E1 has dual functions as an E2 in regulating protein function (for example, through monoubiquitination of MKS1) or targeting them for degradation (for example, polyubiquitination of β-catenin). We therefore asked the question if the co-dependent regulation of MKS1 and UBE2E1 could regulate cellular β-catenin levels. We first confirmed that MKS1 and β-catenin interact (**Figure 6d** and **Supplementary Data 1**) and that sh*Ube2e1* knock-down cells have increased levels of β-catenin (**Figure 6d**), consistent with the up-regulated canonical Wnt/β-catenin signalling that we observed in these cells (**Figure 4d**). We reasoned that, in steady state conditions, UBE2E1 could mediate correct polyubiquitination levels of β-catenin, followed by subsequent targeted degradation, maintaining regulated levels of canonical Wnt signalling. In the event of high levels of the E2, caused by absence of the regulator MKS1, β-catenin is over-polyubiquitated and its levels increase leading to dysregulation of canonical Wnt signalling (**Figure 7**). Consistent with the mutual inhibition of MKS1 and UBE2E1, TUBE pull-down assays confirmed that levels of polyubiquitinated β-catenin increased following *MKS1* knockdown (**Figure 6e**). As expected, levels of polyubiquitinated β-catenin further increased in presence of the catalytically-inactive dominant negative (DN) form of UBE2E1 compared to wild-type (WT) UBE2E1 (**Figure 6e**). We also observed increased levels of phosphorylated β-catenin following loss of MKS1 (**Figure 1a**) and specific localization of phosphorylated β-catenin at the base of the cilium (**Figure 6f**). Levels of phosphorylated β-catenin were also increased at the base of the cilium in *MKS1* knockdown cells, suggesting that this is the cellular location where the phosphorylated form of β-catenin is processed by UBE2E1 for polyubiquitination. These observations indicate that correct β-catenin levels are tightly regulated at the primary cilium by a ciliary-specific E2 (UBE2E1) and a regulatory substrate-adaptor (MKS1).

**Figure 7:**
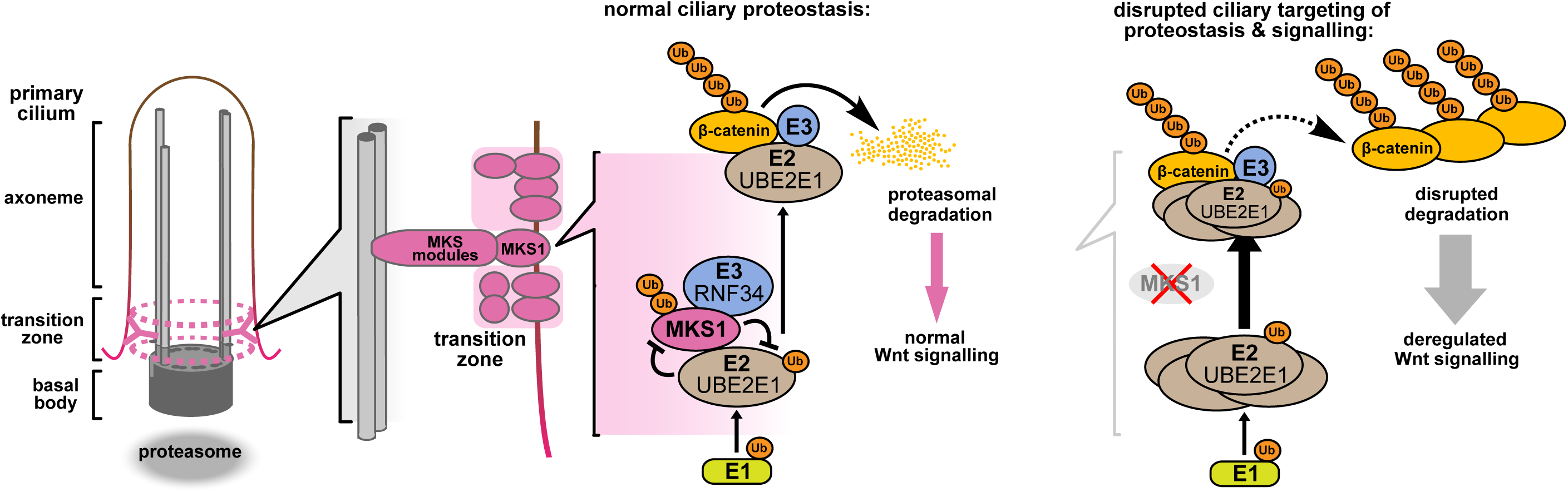
Schematic representation of UPS regulation of MKS1 and β-catenin protein levels at the ciliary apparatus. Protein levels of MKS1 (pink) and UBE2E1 (light brown) are co-dependant through regulation at the base of the cilium. MKS1 localizes to the TZ (dashed pink lines) and is mono-/bi-ubiquitinated by a complex that includes UBE2E1 and RNF34 (blue). MKS1 and UBE2E1 regulate each other, what has an effect on downstream UBE2E1 role in regulation of polyubiquitination of β-catenin (yellow). The correct regulation between these proteins facilitates normal proteasomal function and canonical Wnt signalling (small pink arrow). Both processes are de-regulated following MKS1 mutation of loss (red cross), causing aberrant accumulation of UBE2E1 and polyubiquitinated β-catenin and disrupted tethering to the ciliary apparatus.

## Discussion

A number of studies suggest that the primary cilium or basal body constrains canonical Wnt/β-catenin signalling activity ^4,22,23,39^, and de-regulated, increased signalling is one of the hallmarks of the ciliopathy disease state. Canonical Wnt/β-catenin signalling is aberrantly up-regulated in several ciliopathy animal models, and, in particular, in postnatal cystic kidneys ^40,41^. We have shown previously that homozygous *Mks1*^*-/-*^ mouse embryos also have up-regulated canonical Wnt signalling, reduced numbers of primary cilia and increased proliferation in the cerebellar vermis and kidney ^21^. The mechanistic detail of Wnt signalling de-regulation in ciliopathies remains unclear and controversial. A key question remains whether this ciliary signalling defect is a secondary consequence of cilia loss, or if it is directly and causally related to the loss of function of specific cilium proteins. Several studies support the latter hypothesis, including one study that suggests that jouberin, a component of the TZ/basal body, may modulate Wnt/β-catenin signalling by facilitating nuclear translocation of β-catenin in response to Wnt stimulation ^41^. Regulation of Wnt signalling appears to be also mediated by a functional association of the basal body with the UPS ^29^, through which signalling pathway components such as β-catenin are degraded ^25^. Early studies showed that the basal body and the proteasome can colocalise ^42,43^ and normal, regulated Wnt signalling has been shown to be dependent on the interaction of the basal body protein BBS4 with RPN10, a component of the proteasome ^29^. *Rpgrip1l*^-/-^ knock-out mice have decreased proteasome activity, and a component of the proteasome (Psmd2), was shown to interact with Rpgrip1l ^31^. Furthermore, a number of UPS proteins have been implicated in ciliopathies including TOPORS, an E3 ligase located at the basal body and cilium that is mutated in retinitis pigmentosa (RP) ^44^, and TRIM32, an E3 mutated in Bardet-Biedl syndrome (BBS) ^45^. Interestingly, TRIM32 has been shown to interact with UBE2E1 ^46^. UPS components have also been shown to interact with ciliopathy proteins, such as USP9X with lebercilin ^30^, and the UPS was an enriched biological module that we identified in a whole genome siRNA screen of ciliogenesis ^47^. These observations therefore support a specific role for MKS1 in UPS-mediated proteostasis and signalling regulation.

Here, we demonstrate that loss of MKS1 causes aberrant accumulation of β-catenin (**Figure 1a**) and aberrantly increased proteasome activity (**Figure 1c-d**). The increase in proteasomal activity may be a non-specific response to cellular stress in the absence of MKS1, but our discovery and validation of direct interactions of MKS1 with two proteins (UBE2E1 and RNF34) in the ubiquitination cascade suggest that loss of MKS1 causes a more specific defect. We confirmed biochemical and functional interactions of MKS1 with both UBE2E1 and RNF34 (**Figure 2c, Figure 2e-f, Supplementary Figure 3**), as well as their co-localisation with the basal body (**Figure 3, Supplementary Figure 3**). Loss or dominant negative expression of UBE2E1 mimicked the cellular phenotype of *MKS1* mutants (**Figure 4a-d, g-h**), suggesting a functional interaction between MKS1 and UBE2E1 at the cilium. Using western blotting (**Figure 5d**) we substantiated an inverse correlation between MKS1 and UBE2E1 protein levels in the cell, and using *in vitro* ubiquitination assays (**Figure 6c**) we show that this is due (at least in part) to ubiquitination of MKS1 by UBBE2E1. We suggest that this interaction between MKS1 and UBE2E1 plays a role in regulating Wnt signalling at the base of the primary cilium. In support of this, we demonstrated a functional interaction between UBE2E1, MKS1 and β-catenin (**Figure 6d, e**) and that phosphorylated β-catenin localized at the base of the cilium (**Figure 6f**), presumably prior to UPS processing.

Our data indicates that MKS1 acts as a novel substrate-adaptor that interacts with UPS components and β-catenin, thereby regulating levels of β-catenin through normal degradation during Wnt signalling. MKS1 could mediate the degradation of β-catenin by controlling the stability and the localisation of UBE2E1 at the ciliary apparatus, and perhaps ensuring the correct processing of ubiquitinated β-catenin through close proximity to the proteasome at the ciliary base. This suggestion is supported by the biochemical interaction of another ciliopathy protein, Rpgrip1l, with proteasome proteins and the discrete localization of ubiquitin at the ciliary base ^31^. Catalytically active UBE2E1 regulated ciliogenesis (**Figure 6c**), which implies that UBE2E1-mediated ubiquitylation of substrates such as MKS1 and β-catenin is required for ciliogenesis ^41,42^. In addition, we show for the first time, that MKS1 is polyubiquitinated with non-degradative Lys63-linked chains, which have been shown to have scaffolding roles in other cell signalling networks by bridging together large signalling complexes ^48^. Since MKS1 contains a predicted lipid-binding B9/C2 domain, MKS1 may therefore act as a membrane anchor to ensure the spatial organization and co-ordinated regulation of both the β-catenin destruction complex ^22^ and UPS components at the ciliary apparatus. Loss of MKS1 would lead to the disruption of both the structure and function of the TZ, preventing regulated ciliary signalling and β-catenin degradation (**Figure 7**). In summary, our results indicate that the MKS1-UBE2E1 complex plays a key role in the degradation of β-catenin, which in turn facilitates correct cell function and signalling. Our data provide a mechanistic explanation for Wnt signalling defects in ciliopathies and highlights new potential targets in the UPS for therapeutic intervention.

## Materials and methods

### Informed consent for use of patients in research

Informed consent was obtained from all participating families or patients, with studies approved by the Leeds (East) Research Ethics Committee (REC ref. no. 08/H1306/85) on 4^th^ July 2008.

### Animals

The animal studies described in this paper were carried out under the guidance issued by the Medical Research Council in *Responsibility in the Use of Animals for Medical Research* (July 1993) in accordance with UK Home Office regulations under the Project Licence no. PPL40/3349. B6;129P2-Mks1 ^tm1a(EUCOMM)Wtsi^ heterozygous knock-out mice were derived from a line generated by the Wellcome Trust Sanger Institute and made available from MRC Harwell through the European Mutant Mouse Archive http://www.emmanet.org/ (strain number EM:05429). The *Ub*^*G76V*^*-GFP* line (25) B6.Cg-Tg(*ACTB-Ub\**^*G76V*^*/GFP*)1^Dant/J^ (strain number 008111) was obtained from the Jackson Laboratory, Maine, USA. Genotyping was done by multiplex PCR on DNA extracted from tail tips or the yolk sac of E11.5-E15.5 embryos, or ear biopsies of adult mice. Primer sequences are available on request. Proteasome inhibition treatment of *Mks1* x *Ub*^*G76V*^*-GFP* mice using MG262 was carried out as previously described ^32^.

### Preparation of tissue sections

Mouse embryos or tissue for IF staining were lightly fixed in 0.4% paraformaldehyde, soaked in 30% sucrose/PBS, frozen in OCT embedding medium and cut into 5μm sections on a cryostat. Fresh-frozen sections were left unfixed and processed for immunofluorescent staining by standard techniques.

### Cells

Mouse inner medullary collecting duct (mIMCD3), human retinal pigment epithelium cells immortalized with human telomerase reverse transcriptase (hTERT-RPE1) and immortalized adult retinal pigment epithelium (ARPE19) cells were grown in Dulbecco’s minimum essential medium (DMEM)/Ham’s F12 supplemented with 10% foetal calf serum at 37°C/5% CO_2_. Human embryonic kidney (HEK293) cells were cultured in DMEM with 10% foetal calf serum at 37°C/5% CO_2._ The derivation and culture of mouse embryonic fibroblasts (MEFs) has been described previously ^49^. MEFs were grown in DMEM/Ham’s F12 supplemented with 10% foetal calf serum and 1% penicillin streptomycin at 37°C/5% CO_2_. Fibroblasts from a normal undiseased control, a patient (MKS-562) with a compound heterozygous *MKS1* mutation, and a female patient with a homozygous *ASPM* mutation, were immortalised following transduction with an amphotropic retrovirus encoding the hTERT catalytic subunit of human telomerase, and maintained in Fibroblast Growth Medium (Genlantis Inc. San Diego, CA) supplemented with 0.2 mg/ml geneticin. Patient MKS-562, a compound heterozygote for the *MKS1* mutations [c.472C>T]+[IVS15-7_35del29] causing the predicted nonsense and splice-site mutations [p.R158*]+[p.P470*fs*\*562], has been described previously ^10^. Proteasome inhibition treatment was carried out using 10μM final concentration of the inhibitor dissolved in DMSO for 16 hours (unless otherwise stated). DMSO was used as the vehicle-only negative control.

### Cloning, plasmid constructs and transfections

Human *MKS1* was cloned into the pCMV-cmyc vector as described previously ^50^. The pGEX5X-1-UBE2E1 and pCMV-UBE2E1-FLAG-cmyc constructs have been described previously ^33^. The c.341T>A, p.C131S active site dominant negative (DN) missense mutation was introduced into pCMV-UBE2E1-FLAG-cmyc using the QuickChange mutagenesis kit (Stratagene Inc.) and verified by DNA sequencing. For transfection with plasmids, cells at 80% confluency were transfected using Lipofectamine 2000 (Invitrogen Inc.) according to the manufacturer’s instructions and as described previously ^50^. Cells transfected with plasmids expressing *Ube2e1* shRNA (Origene) were selected for using 0.5μg/ml puromycin for 5 passages. Transfection with Dharmacon ON-TARGET PLUS siRNAs was carried out using Lipofectamine RNAiMAX according to the manufacturer’s instructions and as described previously ^50^. To assess co-dependency of protein levels, 1μg of cmyc-MKS1 was co-transfected with 1, 2.5 and 5μg of FLAG-cmyc-UBE2E1. To investigate if an increased amount of MKS1 would have an effect on UBE2E1 levels, 3μg of cmyc-MKS1 were co-transfected with 1μg FLAG-cmyc-UBE2E1. After 24hr incubation with transfection complexes, cells were treated with 100μg/ml cycloheximide for 4hr. Ubiquitination of cmyc-MKS1 in mIMCD3 cells was assessed after treatment with proteasome inhibitor (MG132 at 10μM) for 3hr.

### Antibodies

The following primary antibodies were used: mouse anti-cmyc clone 9E10, mouse anti-acetylated-α-tubulin clone 6-11B-1, mouse anti-HA (Sigma-Aldrich Co. Ltd.), rabbit anti-GFP (“Living Colors A.v. Peptide Antibody”) and mouse anti-UBE2E1 (BD Biosciences Inc.); rabbit-anti-γ-tubulin and mouse anti-β-actin clone AC-15 (Abcam Ltd.); mouse anti-cyclin D1 clone A-12 (Santa Cruz Biotechnology Inc.); rabbit anti-phospho-β-catenin and rabbit anti-β-catenin (Cell Signalling Technology Inc.); and mouse anti-mono- and polyubiquitinylated conjugates clone FK2 and rabbit anti-20S proteasome α7 subunit (Enzo Life Sciences Inc.). Rabbit anti-MKS1 has been described previously ^51,52^. Secondary antibodies were AlexaFluor488-, and AlexaFluor568-conjugated goat anti-mouse IgG and goat anti-rabbit IgG (Molecular Probes Inc.) and HRP-conjugated goat anti-mouse immunoglobulins and goat anti-rabbit immunoglobulins (Dako Inc.).

### Immunofluorescence and confocal microscopy

Cells were seeded at 1.5 × 10^5^ cells/well on glass coverslips in six-well plates, 24 hours before transfection and 48-96 hours before fixation. Cells were fixed in ice-cold methanol (5 minutes at 4°C) or 2% paraformaldehyde (20 minutes at room temperature). Permeabilization, blocking methods and immunofluorescence staining were essentially as described previously ^53^. Confocal images were obtained using a Nikon Eclipse TE2000-E system, controlled and processed by EZ-C1 3.50 (Nikon Inc.) software. Images were assembled using Adobe Photoshop CS3 and Adobe Illustrator CS2.

### Yeast 2-hybrid screening

The B9/C2 domain of human *MKS1* (amino acids 144-470; **Figure 4a**) was cloned into the Gal4 vector pB27 and screened against a human fetal brain RP1 prey cDNA library. Yeast-2-hybrid screens were performed by Hybrigenics SA as described previously ^50^. Confirmatory “1-to-1” pairwise assays for selected interactants were performed with the MatchMaker Two-Hybrid System 3 (Clontech Inc.)

### GST fusion protein purification

GST-UBE2E1 fusion protein was prepared essentially as described previously ^33^, with protein expression induced at 20°C using 0.2mM IPTG for 4 hr.

### Proteasome activity assays

Crude proteasomal fractions were prepared from cells ^54^ and incubated with the 20S fluorophore substrate Suc-LLVY-AMC (Enzo Life Sciences Inc.). Fluorescence of each proteasomal preparation was measured on a Mithras LB940 (Berthold Technologies Inc.) fluorimeter and adjusted against a calibration factor calculated from a standard curve to give activity measurements in pmol AMC release/µg cell lysate/hour. Treatment of cells with 10μM of the proteasome inhibitors MG-132 or c-lactacystin-β-lactone were positive controls for the assay. Results reported are from at least five independent biological replicates.

### Canonical Wnt activity (SUPER-TOPFlash) luciferase assays

For luciferase assays of canonical Wnt activity, we grew cells in 12-well plates and co-transfected with 0.5 μg SUPER-TOPFlash firefly luciferase construct^55^ (or FOPFlash, as a negative control); 0.5 μg of expression constructs (pCMV-cmyc-MKS1, or empty pCMV-cmyc vector); and 0.05 μg of pRL-TK (Promega Corp; *Renilla* luciferase construct used as an internal control reporter). We obtained Wnt3a- or Wnt5a-conditioned media from stably-transfected L cells with Wnt3a or Wnt5a expression vectors (ATCC). Control media was from untransfected L cells. Activities from firefly and *Renilla* luciferases were assayed with the Dual-Luciferase Reporter Assay system (Promega Corp.) on a Mithras LB940 (Berthold Technologies Inc.) fluorimeter. Minimal responses were noted with co-expression of the FOP Flash negative control reporter construct. Raw readings were normalized with *Renilla* luciferase values. Results reported are from at least four independent biological replicates.

### Ubiquitination assays

To assess *in vitro* ubiquitination, we used a ubiquitination kit (Enzo Life Sciences, Inc.) according to the manufacturer’s protocol, supplemented with MKS1-GST (Proteintech Group, Inc.) and RNF34-HIS (Novus Biologicals) fusion proteins in a total volume of 30μl. Samples were incubated for 1.5hr at 37°C followed by SDS-PAGE and Western blotting.

### TUBE assays

Agarose-bound TUBE were used as recommended by the manufacturer (LifeSensors, Malvern, PA, USA). mIMCD3 cells were transiently transfected with cmyc-MKS1 and treated with proteasome inhibitor (MG132 at 10μM) 2hr before harvesting. Lysis buffer was based on RIPA supplemented with 50mM Tris-HCl pH7.5, 0.15M NaCl, 1mM EDTA, 1% NP40, 10% glycerol, DUB inhibitors (50μM PR619 and 5mM 1,10-phenanthroline) and protease inhibitors. BRISC was used as Lys63 deubiquitinating enzyme. Samples were run on SDS-PAGE followed by Western blotting using standard protocol.

### Co-immunoprecipitation and mass spectrometry

Whole cell extracts (WCE) were prepared and co-IP performed essentially as described previously ^56^. Co-IPs used either 5 µg affinity-purified mouse monoclonals (MAbs), or 5-10 µg purified IgG fractions from rabbit polyclonal antisera, coupled to protein G- and/or protein A-sepharose beads (GE Healthcare UK Ltd.). Proteins were eluted from beads with 0.2M glycine HCl pH2.5. Samples were neutralised by addition of 0.1 volume 1M Tris HCl ph8.5. After elution, proteins were precipitated with chloroform and methanol and subjected to in-solution tryptic cleavage as described previously ^57^. LC-MS/MS analysis was performed on Ultimate3000 nano RSLC systems (Thermo Scientific) coupled to a Orbitrap Fusion Tribrid mass spectrometer (Thermo Scientific) by a nano spray ion source ^58^. Mascot (Matrix Science, Version 2.5.1) was used to search the raw spectra against the human SwissProt database for identification of proteins. The Mascot results were verified by Scaffold (version Scaffold_4.8.8, Proteome Software Inc., Portland, OR, USA) to validate MS/MS-based peptide and protein identifications.

### Western blotting

Soluble protein was analysed by SDS-PAGE using 4-12% Bis-Tris acrylamide gradient gels and western blotting was performed according to standard protocols using either rabbit polyclonal antisera (final dilutions of x200-1000) or MAbs (x1000-5000). Appropriate HRP-conjugated secondary antibodies (Dako Inc.) were used (final dilutions of x10000-25000) for detection by the enhanced chemiluminescence “Femto West” western blotting detection system (Pierce Inc.). Chemiluminescence was detected using a BioRad ChemiDoc MP Imaging System and Image Lab software. Volumetric analysis of immunoblot bands was performed using Image Lab software (Bio Rad).

### Statistical analyses

Normal distribution of data (for SUPER-TOPFlash assays, proteasome activity assays, cilia length measurements) was confirmed using the Kolmogorov-Smirnov test (GraphPad Software). Paired or unparied comparisons were analysed with either Student’s two-tailed t-test or χ^2^ tests using InStat (GraphPad Software). Results reported are from at least three independent biological replicates.

## Acknowledgements

This paper is dedicated to the memory of P. Robinson, our valued collaborator, colleague and friend. We are very grateful to D. Evans, J. Bilton, C. McCartney and M. Reay for technical support. We thank E. Pitt for hTERT immortalization of MKS1 patient fibroblasts. We thank A. Monk, K. Passam and T. Simpson of Nikon UK Ltd. for technical support and advice on confocal microscopy. We are grateful to R. T. Moon (University of Washington) for the SUPER-TOPFlash and FOPFlash constructs, and S. Kang (Korea University) for the UBE2E1 constructs. The anti-MKS1 antibody was a gift from N. Katsanis, Duke University Medical Center. We acknowledge funding from the UK Medical Research Council (Doctoral Training Award for GW). The research also received funding from the European Community’s Seventh Framework Programme FP7/2009 under grant agreement no: 241955 SYSCILIA. KS was funded by Wellcome Trust Institutional Strategic Support Funding to University of Leeds, and GW was funded by a Wellcome Trust Seed Award in Science (204378/Z/16/Z).

## Author contributions

G.W. and C.A.J. contributed equally to this manuscript by conceiving of the study, supervising the research and analyzing data. K.S. and G.W. performed cell biology and biochemical experiments; K.B. and M.U. performed, analyzed and supervised mass spectroscopy experiments. C.V.L. and M.A. validated biochemical interactions and interpreted data. P.A.R. and E.Z. provided reagents and interpreted data. All authors contributed to drafting the manuscript and reviewed the final submission.

## Supplementary Figure legends

**Supplementary Figure 1: Characterisation of *MKS1*-mutated human patient fibroblasts (a)** RT-PCR amplicons of exons 4-6 and 15-17 from cDNA of healthy control and MKS1 patient fibroblasts, compound heterozygote for the *MKS1* mutations [c.472C>T]+[IVS15-7_35del29] causing the predicted nonsense and splice-site mutations [p.R158*]+[p.P470*fs*\*562]. Additional smaller PCR products in MKS1 patient corresponds to skipping of exon 5 and exon 16, confirmed by Sanger sequencing, due to the frameshift mutation affecting splicing. **(b)** Immunoblot showing loss of MKS1 protein in *MKS1*-mutated patient fibroblasts compared to healthy controls; loading control is β-actin. **(c)** IF microscopy images of wild-type control and *MKS1*-mutated fibroblasts showing loss of cilia and disorganization of cytoskeleton in patient cells. Bar graphs quantify reductions in incidence and length of cilia in patient cells. Statistical significance of pairwise comparisons with control for three independent biological replicates are shown (*** *p* < 0.001, **** *p* < 0.0001; paired two-tailed Student t-test; error bars indicate s.e.m.) **(d)** IF microscopy images showing loss of MKS1 ciliary localization in *MKS1*-mutated patient fibroblasts compared to wild-type control fibroblasts (indicated by arrowheads).

**Supplementary Figure 2: *In vivo* loss of MKS1 causes deregulated ubiquitin-proteasome processing. (a)** Proteasome defects (GFP; green) in *Mks1*^*-/-*^ x *Ub*^*G76V*^*-GFP* E12.5 embryonic heart and liver, as indicated. Scale bars = 50 μm. **(b)** Left panel: accumulation of active β-catenin (red) and GFP-tagged ubiquitin (green)in the neuroepithelium of E12.5 *Mks1*^*-/-*^*xUb*^*G76V*^*GFP* embryonic neocortex. Middle panel: loss of primary cilia (stained for acetylated α-tubulin, red) on the neuorepithelial cells lining the fourth ventricle in mutant animals. Right panel: loss of basal body localisation of UBE2E1 (red; indicated by arrowheads) in mutant animals, accompanied by accumulation GFP-tagged ubiquitin. The asterisk indicates a periventricular heterotopia. Scale bar = 10μm. White frames indicate magnified regions displayed in insets (showing only the red and green channels for clarity).

**Supplementary Figure 3: The E3 ubiquitin ligase RNF34 interacts with MKS1 and co-localizes at the basal body. (a)** Co-immunoprecipitation of 30 and 40 kDa isoforms of endogenous RNF34 by rabbit polyclonal anti-MKS1, but not by an irrelevant antibody or the pre-immune. **(b)** Co-localization of RNF34 (red) with MKS1 (green) at the basal body (arrowheads) in mIMCD3 cells. Scale bar = 10μm.

**Supplementary Figure 4: Validation of *Ube2e1* siRNA knockdown in mIMCD3 cells and effect of shRNA *Ube2e1* knockdowns on MKS1 interacting proteins under different conditions of ciliogenesis. (a)** Left panels: mouse *Ube2e1* pooled siRNA duplexes (si*Ube2e1*) prevent ciliogenesis compared to scrambled (siScr) control. Primary cilia and basal bodies are visualized by staining for acetylated α-tubulin (red) and γ-tubulin (green). Scale bar = 10 μm. Right panel: immunoblot to confirm reduction of Ube2e1 protein levels following knockdown in mIMCD3 cells using individual siRNA duplexes (#1 to #3) and pooled duplexes, with knockdown efficiency indicated normalized to β-actin loading control levels. **(b)** Scatter plot representing log_2_ fold-difference values for peptide ratios shScr: sh*Ube2e1* in mIMCD3 knockdown cells versus -log_10_ *p* χ^2^ tests for shUbe2e1: shScr rations across different conditions of ciliogenesis. These conditions comprise: proliferating cells grown in media containing serum (red); ciliated cells grown in the absence of serum (grey); and cells undergoing ciliary resorption grown in the absence of serum followed by 2hr incubation in media with serum (gold). The proteins indicated in the plot are members of the enriched biological process GO term “actin filament capping” term which includes known interactants of ciliopathy proteins such as filamin A and B (Flna and Flnb). Full data-sets for this plot are available in **Supplementary Dataset 1**.

**Supplementary Dataset 1: Mass spectrometry results for MKS1 pull-downs from mIMCD3 cells across different conditions of ciliogenesis.** The data-set lists proteins (identified by ≥3 unique peptide counts) pulled-down by anti-MKS1 immunoprecipitations, under different conditions of ciliogenesis (proliferating cells, ciliated cells, cells undergoing ciliary resorption). Columns A and B: protein and gene name. Column C: protein accession number. Columns G to L: peptide counts identified by LC-MS/MS mass spectrometry (columns F and M indicating non-specific peptide counts following BSA washes), with counts heat-mapped red (high) to green (low). Column N: χ^2^ tests of peptide counts for shScr compared to sh*Ube2e1* knockdown cells, across different conditions of ciliogenesis (red highlighted cells indicate χ^2^ test *p*<0.05). Columns Q to S: shScr: sh*Ube2e1* peptide count ratios (derived from columns G to L), with values <1 indicating decreased peptide counts following sh*Ube2e1* knockdown. Columns U to Z: indicate if a particular protein was identified in a significantly enriched biological process under the indicated GO terms (row 2).

