## Supplementary figures and images for "Regulation of canonical Wnt signalling by the ciliopathy protein MKS1 and the E2 ubiquitin-conjugating enzyme UBE2E1"

### Supplementary figures

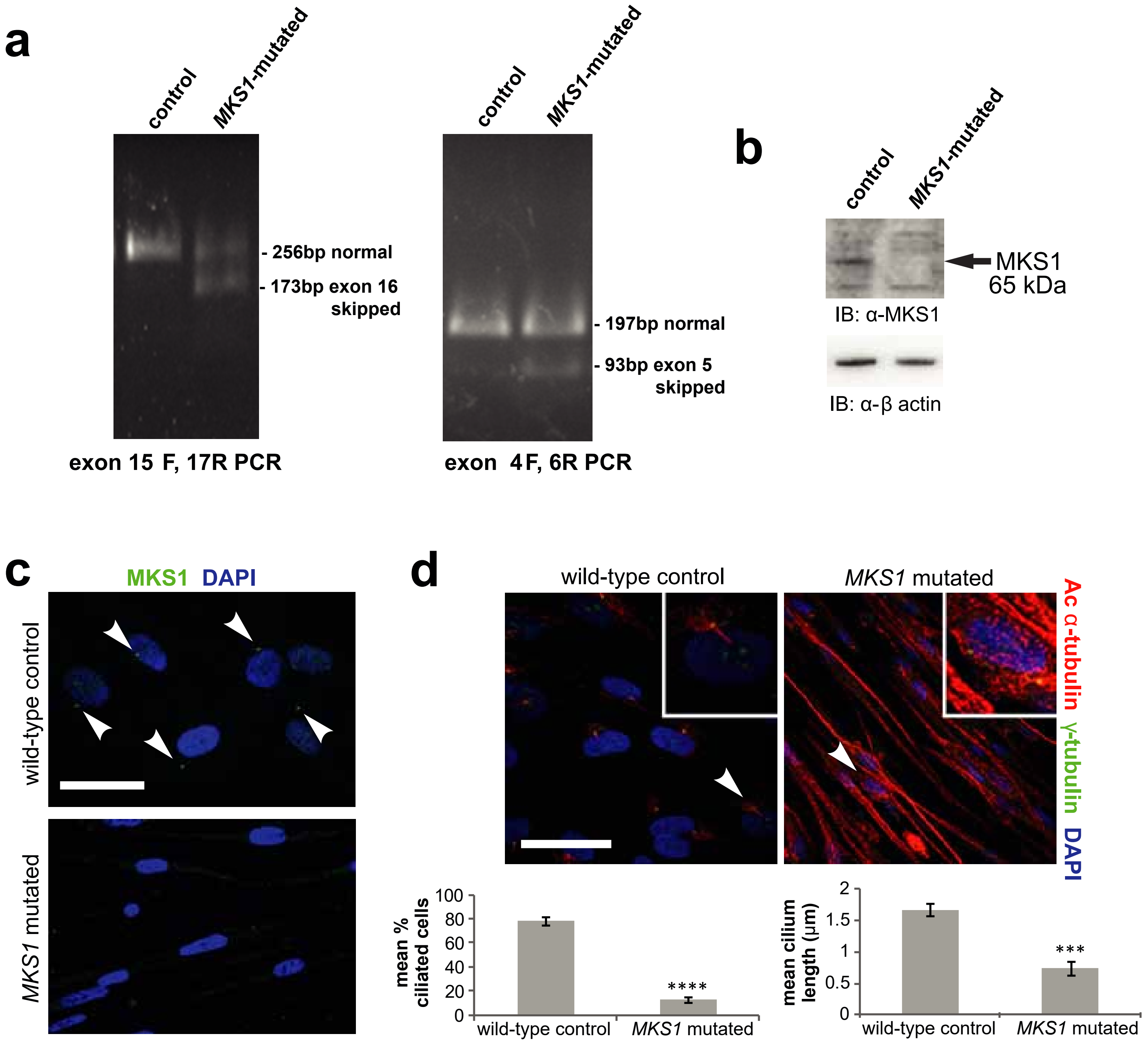

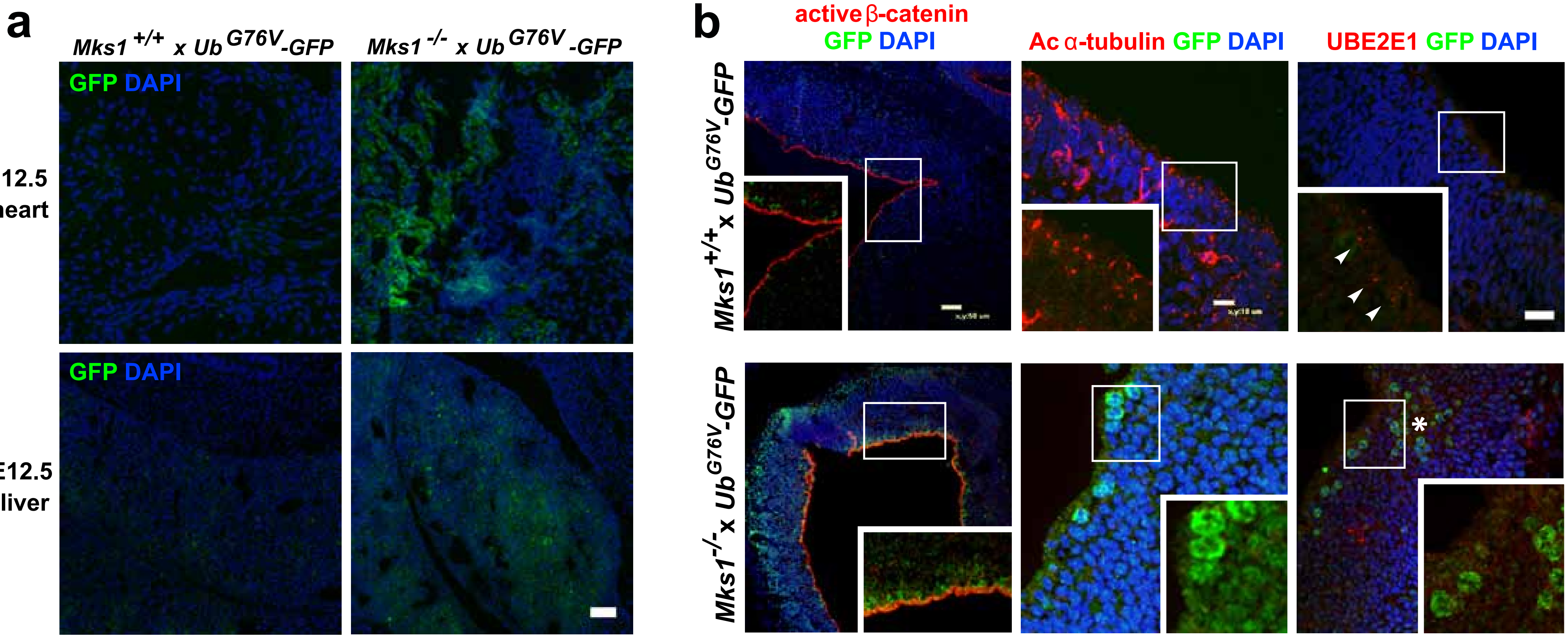

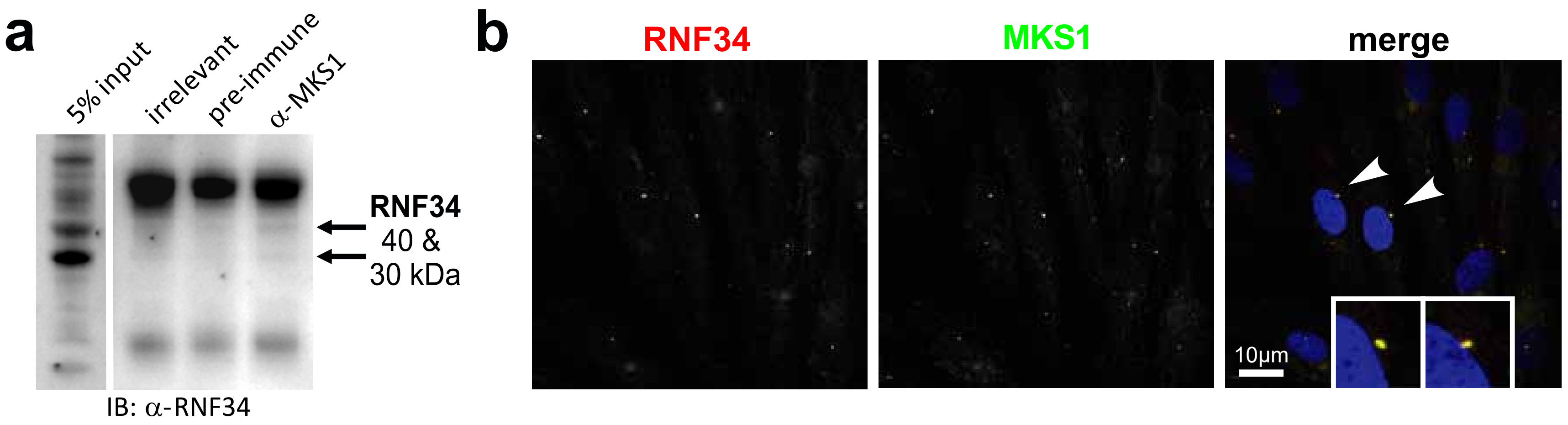

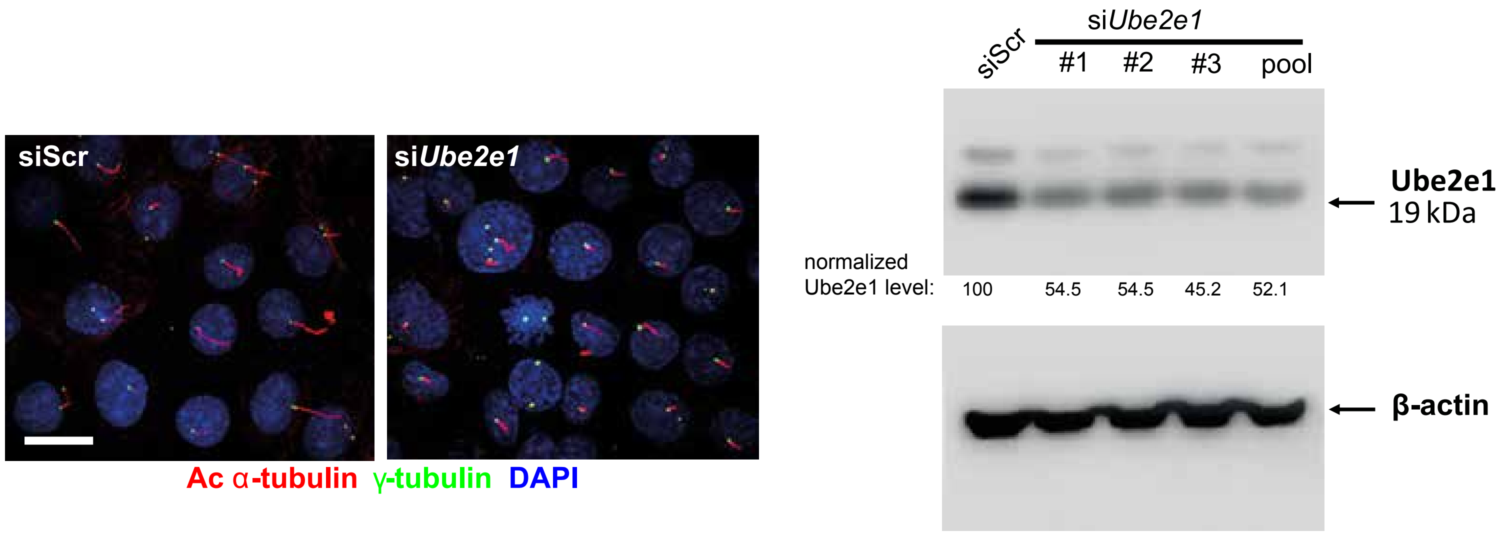

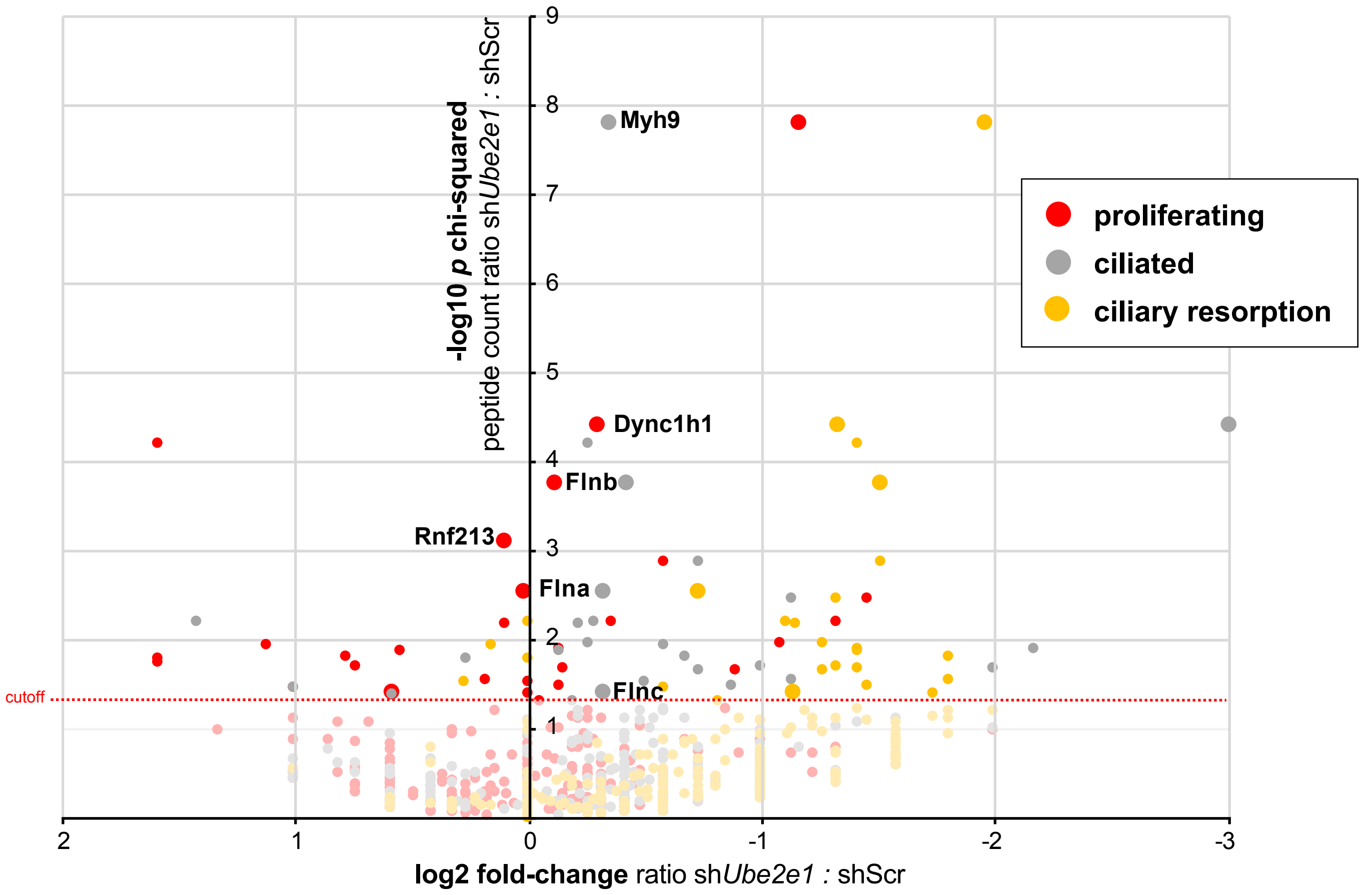
